## Supplementary Material for "A deep learning-based stripe self-correction method for stitched microscopic images"

#### Table of contents

**Supplementary Note 1: Implementation details of proximity sampling strategy**
**Supplementary Note 2: Procedure of artifact and stripe synthesis**
**Supplementary Note 3: Reasons and limitations of using deep learning comparison methods**
**Supplementary Note 4: Training and testing details**
**Supplementary Note 5: Ablation study**
**Supplementary Note 6: Brief introduction to the comparison methods**

**Supplementary Figure 1: Representative sampling cases of stripe and artifact**
**Supplementary Figure 2: Other representative stripe correction results**
**Supplementary Figure 3: Detailed correction results of imprecise prior stitched information**
**Supplementary Figure 4: Pipeline of the stripe and artifact synthesis**
**Supplementary Figure 5: Restoration results of real artifacts in the MPM images**
**Supplementary Figure 6: Cell counting results on bubble artifact and stripe images**
**Supplementary Figure 7: Accuracy of automatic counting compared with manual counting**
**Supplementary Figure 8: Visualization of network architecture**

**Supplementary Table 1: Requirements of different methods for training images**
**Supplementary Table 2: Comparison of the microscopic datasets**

### Supplementary Notes

##### Supplementary Note 1: Implementation details of proximity sampling strategy

For the stitched images with stripes, first of all, the partition of tiles is roughly annotated by hand. According to the dimension of the tiles, we empirically determine the dimension of sampling patches. For the label-free MPM image of breast cancer, the resolution of the tile is approximately 512×512 pixels while the resolution of the sampling patches is set as 256×256 pixels. For the liver cancer image, the width of grid stripe is less than 128 pixels, so we sample patches with the dimension 128×128 pixels.

In order to train SSCOR, each normal patch needs to pair with an anomaly patch. In our proposed proximity sampling strategy, we allow each anomaly patch to pair with a normal patch in proximity. The representative sampling examples with regards to stripe and artifacts are shown in Supplementary Fig. 1, the green boxes refer to the sampled anomaly patches, and the blue boxes are the corresponding normal ones. Notably, the blue dashed boxes denote the sub-normal patches that should not to be sampled as normal patches. In Supplementary Fig. 1a, three typical anomaly patches are located on the horizontal and vertical stripes as well as the junction of stripes, respectively, while their corresponding normal patches are sampled nearby. On the other hand, for the images with special artifacts, the adopted sampling strategy is slightly different. In Supplementary Fig. 1b, we display an exemplar image containing not only stripes but also out-of-focus artifacts. In the left area with artifacts, we densely sample the anomaly patches on stripes and sample the corresponding normal patches from the proximity area outside the artifacts.

**Supplementary Note 2: Procedure of the artifact and stripe synthesis**

To precisely assess the restoration quality for the images with three types of artifacts, the clean SRS images are employed as the base images and ground truth, and different artifact masks are combined with them to mimic the images contaminated by artifacts. As follows, the details of generating stripes and artifacts are explained, respectively.

In practice, the stripes and special artifacts may exist in the stitched images at the same time. The process of various artifact synthesis can be formally defined as follows. Specifically, for the stripe, bubble, and out-of-focus artifacts, the synthesis process can be formulated as:

$$I_S = I_R * (1-\alpha) * I_M, \tag{1}$$

where the synthetic mask $I_M$, each pixel of which has an intensity value between zero and one, is combined with the region of raw image $I_R$ to acquire the synthetic result $I_S$. $\alpha$ denotes the balancing factor for the masking impact of stripe and the three artifacts. The larger the value, the greater the effect of the mask, and vice versa. By altering the factor, diverse synthetic images can be obtained.

**Stripe artifacts.** To combine non-uniform stripes with stripe-free stitched images, the mean intensity of all the raw tiles from a stitched MPM image is calculated and normalized, which serves as shading-mask $I_M$. Then, we use each tile of a clean SRS image as $I_R$, and combine it with the mask

$I_M$ to produce the synthesized image $I_S$. By adjusting the balancing factor $\alpha$ in Equation (1), the diverse shading-masks $I_M$ are acquired. Finally, the non-uniform stripe image is generated by stitching these synthesized tiles $I_S$.

**Bubble-like artifacts.** The bubble-like mask $I_M$ is empirically designed to maximally imitate the real bubble-like artifacts in MPM images (Supplementary Fig. 5). In specific, to simulate the non-uniform pattern inside bubble artifacts, the intensity attenuation of the mask $I_M$ is manipulated by a Gaussian filter that smooths the pixel values around each center pixel. Thereafter, the mask $I_M$ is combined with a random region $I_R$ of the clean SRS image.

**Out-of-focus artifacts.** According to previous imaging experience and the real out-of-focus images, the out-of-focus artifacts commonly exists in the four corners of stitched images. Therefore, we empirically generate the mask of out-of-focus artifact $I_M$. The intensity of $I_M$ is set to be faded in periphery, modeled by a Gaussian filter, to imitate the uneven out-of-focus affect. Then, we use the corner of clean SRS image as $I_R$, which will combine with $I_M$ to produce the synthesized image $I_S$.

**Scanning fringe artifacts.** The nonlinear optical images often exhibit strong scanning fringe artifact (SFA) resulting from the fast galvo-resonant (GR) scanning system[1]. According to the reference that appears SFA[1], we observe that SFA often occurs in the blue channel with weak signal areas, due to the low efficiency of nonlinear process. Thus, we employ the blue channel of human ovarian carcinomas images as synthetic mask $I_M$ to generate scanning fringe image $I_S$. The synthesis process can be formulated as:

$$I_S = I_R + \beta * I_M * 255, \tag{2}$$

where $\beta$ refers to the balancing factor similar to $\alpha$. In order to simulate SFA realistically, we define it as additive noise, and simply integrate $I_M$ and $I_R$ through addition operation. We add the mask $I_M$

to the clean SRS images $I_R$ and thus synthesize scanning fringe artifacts on the partial region of the raw image.

**Supplementary Note 3: Reasons and limitations of using deep learning comparison methods**

In the artifact removal experiments, we employ several deep learning-based unsupervised methods for comparison, according to the unsupervised learning settings of our task. As follows, we explain the reason of employing these methods and their respective limitations for our task.

**ZeroDCE**[2] was used in the experiments for removing the out-of-focus artifacts. Since the out-of-focus regions have lower intensity than the other regions in the stitched images, it can be removed by enhancing the brightness of the artifact regions to some extent. We tried several state-of-the-art unsupervised low-light enhancement methods including RUAS[3], SCI[4] and ZeroDCE[2] to explore this feasibility. Among these methods, ZeroDCE achieves the optimal performance. However, based on our observation, there is a large domain gap between the natural image domain and the microscopic image domain in texture contrast and resolution, which makes it difficult to adapt these methods on the task of out-of-focus artifact removal for the microscopic images.

**Mask-ShadowGAN**[5] was involved in the experiments for removing the bubble artifacts. Since the bubble artifacts appear to be similar to the soft shadow in natural images, we employ the state-of-the-art unsupervised shadow removal methods, Mask-ShadowGAN[5] and LG-ShadowNet[6], to remove bubble artifacts. Mask-ShadowGAN shows respectable performance. But, it treats shadow in the form of binary mask, so it excels at handing hard (uniform) shadow rather than soft (non-uniform) shadow. Thus, it does not perform well on the task of bubble artifact removal.

**Neighbor2Neighbor**[7] served as a representative image denoising method in the experiments for removing the scanning fringe artifacts. SFA can be considered as a special type of noise from the imaging system. Thus, we use the off-the-shelf image de-noising method, Neighbor2Neighbor[7] and Self2Self[8], to deal with the noise in scanning fringe artifacts. However, the noise pattern of SFA is significantly different from natural noises, so they fail to recover the original tissue signal from SFA.

#### Supplementary Note 4: Training and testing details

For training SSCOR, we adopt the Adam solver[9] with a learning rate of 0.0002 and the exponential decay rates for the first and second moment estimates that are set to 0.5 and 0.999, respectively. During training, the number of epochs is 200, the learning rate is fixed in the first 100 epochs, and linearly decay to zero over the next 100 epochs. On the inference stage, for the entire stitched image, we densely sample patches following the sliding-window strategy. For the label-free MPM image of breast cancer, with the patch size 256 pixels, the step size of sliding-window sampling is set as 100 pixels. As for the grid stripe MPM images, the stripe only exists in the grid regions, and the stripe size is smaller than 128 pixels. So we set 128 pixels as the patch size and only restore the stripe regions.

#### Supplementary Note 5: Ablation study

Ablation studies were conducted on the proposed schemes, including proximity sampling, reciprocal training, and local-to-global correction, in order to demonstrate the reason for the model design.

##### Proximity sampling

To study the effectiveness of proximity sampling, we compare it with central sampling and random sampling strategies. For all of these sampling strategies, the patches on stripes are regarded as anomaly patches. The main difference rests in the way of sampling normal patches. The central sampling strategy utilizes the central patch of the tile as the normal patch corresponding to the closest anomaly patch, while the random sampling strategy randomly selects the patches from the regions off stripes over the entire stitched image. As illustrated in Supplementary Fig. 9, proximity sample strategy shows superior performance over the other two strategies in the stripe correction and signal restoration.

Next, the dimension of the sampled patches is studied, which depends on the width of stripe and the size of the image tiles. Empirically, it should be smaller than image tile and slightly wider than stripes, in order to enable SSCOR sample the proper normal patches. We compare the stripe removal results with different sizes of patch in Supplementary Fig. 10. Considering the stripe size in most stitched images, we empirically set patch size as 256 as the default setting which obtains the better de-stripe results.

**Adversarial reciprocal-training**

In the adversarial reciprocal training scheme of SSCOR, the stripe synthesis network adds the shadings to the corrected patches in order to preserve image content, since the consistency between the original patches and the synthesized ones serves as effective constrain for stripe correction. We conduct visual comparison experiments to verify the performance with and without stripe synthesis network. As shown in Supplementary Fig. 11, without stripe synthesis network, there still remains the traces of stripes in the corrected image.

**Local-to-global Correction**

As the final stage of SSCOR workflow, local-to-global correction consists of two steps, *i.e.,* local correction and global merging. First, the whole-slide images are partitioned into overlapping local patches in the sliding-window manner and fed into the stripe correction network to obtain corrected patches, which is the local correction step. Next, all the corrected local patches will be merged into a whole image with the same resolution as the original stitched image, which is the global merging step.

To verify our overlapping local correction strategy, we compared it with the other two strategies, stripe-only local correction and non-overlapping local correction, on an MPM image with horizontal stripes and out-of-focus artifacts, where the out-of-focus artifacts exist in the upper left corner (Supplementary Fig. 12). As observed, stripe-only local correction is able to suppress the shading of stripes, but cannot completely correct the out-of-focus areas beyond the stripes. As for non-overlapping local correction, the corrected result may exist stripe residues. To sum up, overlapping local correction obtains the most significant correction results (Supplementary Fig. 12).

**Supplementary Note 6: Brief introduction to the comparison methods**

The comparison methods used in the experiments are introduced in the following.

As a retrospective method, CIDRE[10] operates on a set of image tiles corrupted by illumination distortion that are acquired under consistent conditions. It estimates shading distortion parameters using regularized energy minimization to correct intensity distributions.

BaSiC[11] is another retrospective method for background and shading correction of image tiles, based on a sparse and low-rank decomposition. It imposes sparse constraints on the Fourier-transformed shading model to enforce a good shading correction.

ZEN[12] is a commercial software from Zeiss and it performs shading correction based on a shading reference image generated from image tiles. ZEN requires a raw image with meta information. Moreover, it demands over 300 tiles per image for a good reference image.

ZeroDCE[2] proposes a light-weight deep network for low-light image enhancement. It coverts this task into an image-specific curve estimation problem, which uses image as input, curve as output. The curves perform pixel-wise adjustment on the input images to get enhanced images. It can be trained end-to-end without any reference image by setting a series of non-reference losses, which implicitly evaluate the quality of output image.

Mask-ShadowGAN[5] presents a mask-guided generative adversarial network for shadow removal from unpaired training data. It models the relationship between shadow and shadow-free images by their difference, and uses the shadow mask (binary image) to represent the difference. The network learns to generate a shadow-free image and then produces a shadow image as guided by the shadow mask. By formulating the consistency constraints to obtain shadow images and learns to remove shadows.

Neighbor2Neighbor[7] is a self-supervised image denoising framework with only noisy images. This approach generates sub-sampled paired images by random neighbor sub-samplers from noisy images. Denoise one of the sub-sample image, and construct a reconstruction loss with the other one. Then denoise the original noisy image, and derive sub-sampled pair by using the same neighbor sub-sampler to calculate the regularization term.

### Supplementary Figures

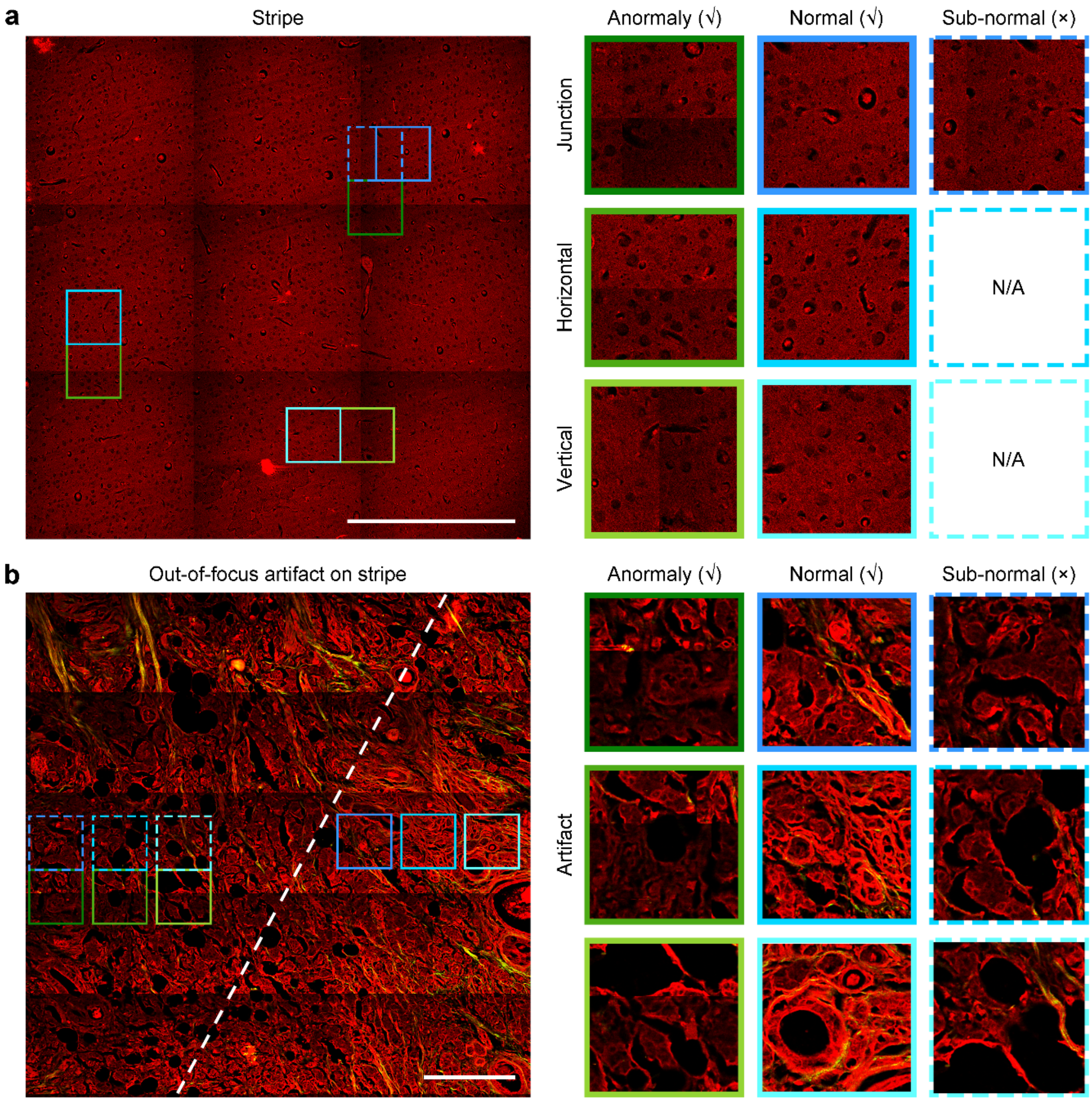


**Supplementary Figure 1. Representative sampling examples with regards to stripe and artifacts. a** In the given stitched stripe image, the anomaly patches (green boxes) on stripes and the corresponding normal patches (blue boxes) in proximity are sampled and paired up for training. In concrete, the anomaly patches on the horizontal (or vertical) stripes pair with the nearest normal patches, i.e., the ones off the stripes along the vertical (or horizontal) direction. Note that, for the anomaly patch on the junction of the vertical and horizontal stripes, the nearest patches off the junction area along the vertical or horizontal direction (blue dashed boxes) are also affected by the stripes, namely sub-normal patches, and thereby they cannot be treated as normal patches for training. Thus, for the anomaly patches resting on the junction, the nearest diagonal patches are chosen as the corresponding normal patch, instead of choosing the sub-normal patches. **b** For the image with both stripes and special artifacts (in addition to the stripes covering the entire figure, it contains the out-of-focus artifacts on the left side of the dotted line), the sampled anomaly patches (green boxes) from artifacts and the corresponding normal patches (blue boxes) constitute the unpaired training data. The nearest patches off the stripes along the vertical (horizontal) direction affected by the artifacts are also considered as sub-normal patches (blue dashed boxes). Likewise, these sub-normal patches cannot be used in training. Therefore, SSCOR samples the patches from the regions with out-of-focus artifacts as anomaly patches, meanwhile sampling the normal patches in the remaining areas without artifact. Scale bars in **a** and **b**: 200 μm.

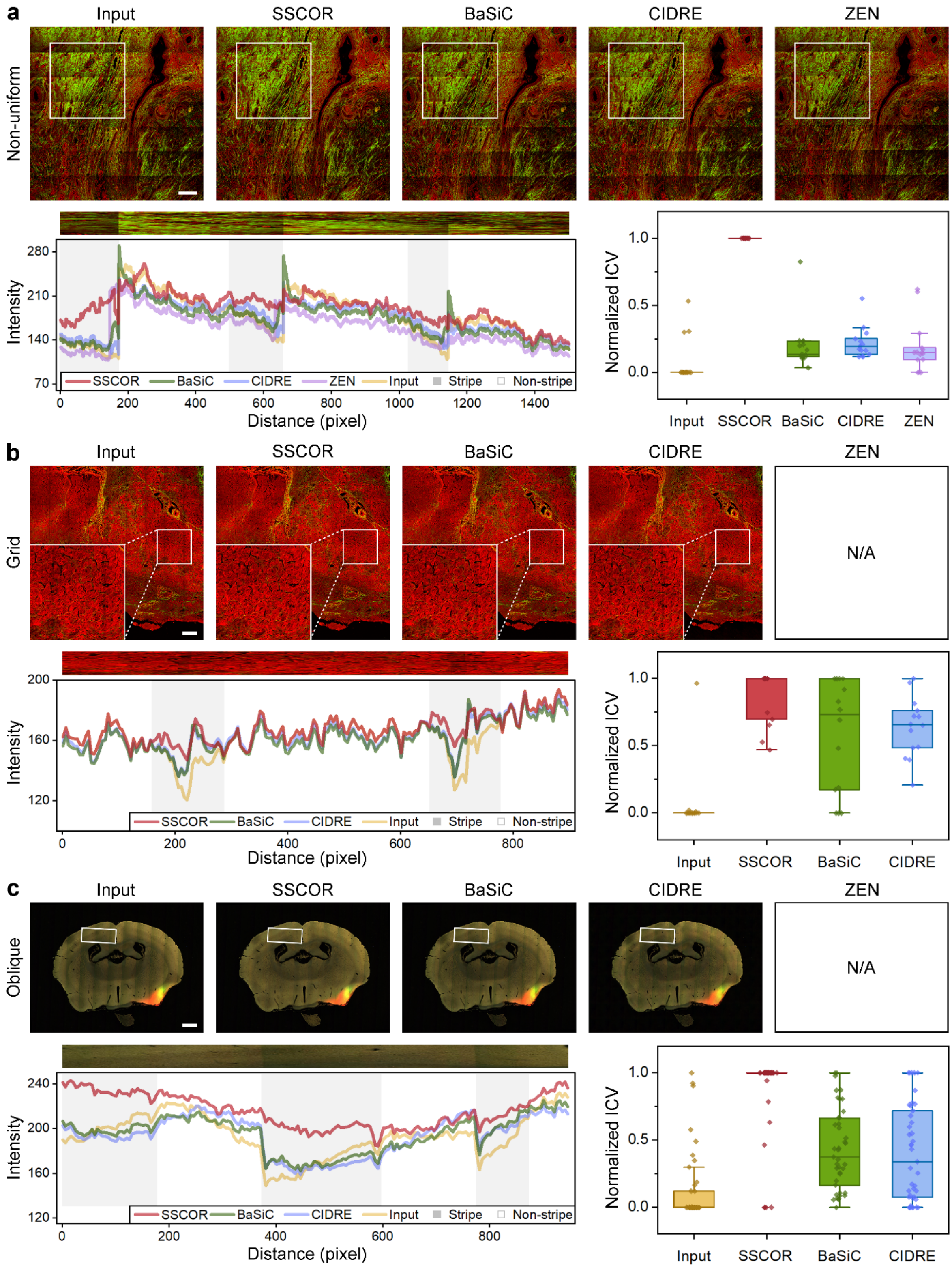


**Supplementary Figure 2. Other representative stripe correction results. a, b,** and **c** Representative correction results and the corresponding quantitative analysis of non-uniform stripe (label-free multiphoton image of breast cancer), grid stripe (label-free multiphoton image of liver cancer), and oblique stripe (labeled fluorescence image of mouse brain[13]). SSCOR performs the state-of-the-art performance for stripe correction compared with the comparison approaches. The Zeiss correction method is not available because there are no raw files for the grid and oblique stripe images. Scale bars in **a** and **b**: 200 μm. Scale bars in **c**: 1 mm.

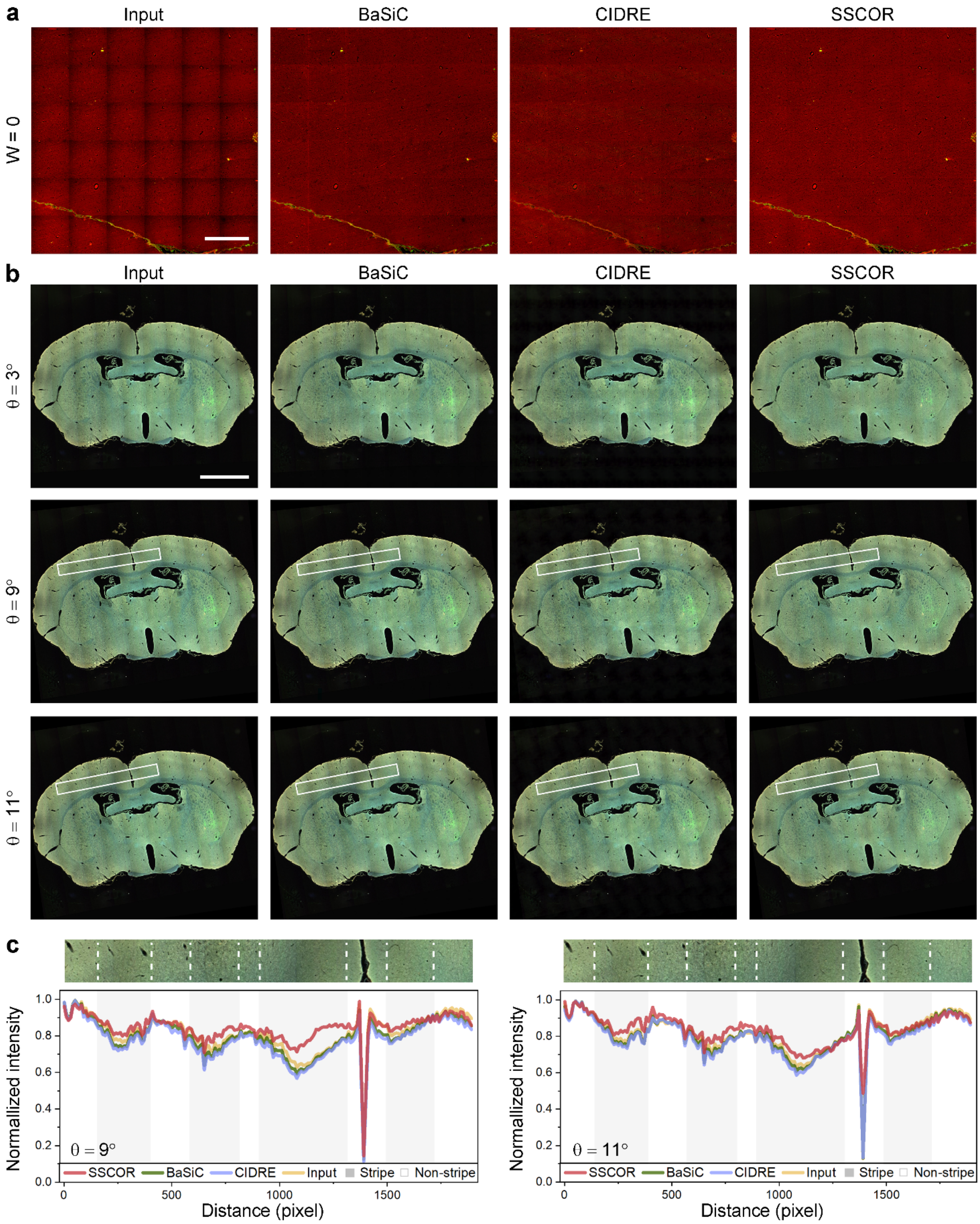


**Supplementary Figure 3. Detailed correction results of imprecise prior stitched information. a** The comparative results of CIDRE[10] and BaSiC[11] under the ROI cropping conditions. Scale bar: 1 mm. **b** The corrected results under the condition of greater image rotation. The intensity profiles within the rectangular regions quantitatively demonstrate that SSCOR has a better correction effect than other comparison methods when the angle of oblique stripes (θ) is no more than 11 degrees. Scale bar: 2 mm.

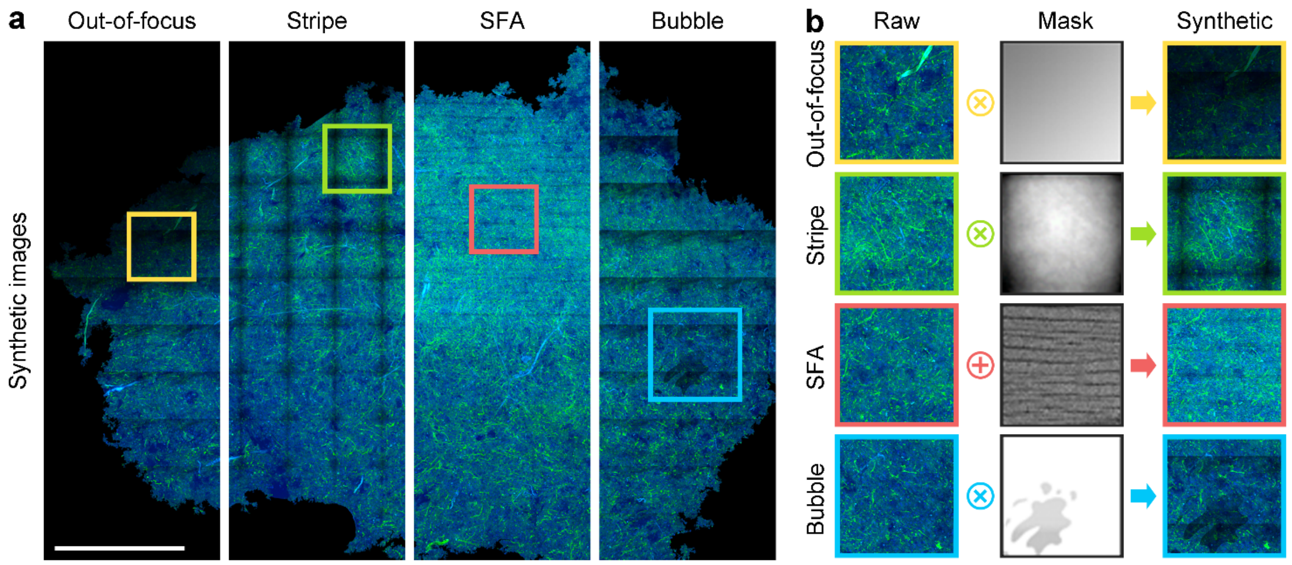


**Supplementary Figure 4. Pipeline of the stripe and artifact synthesis. a** The synthetic images with non-uniform stripe and artifacts (bubble-like, out-of-focus, and scanning fringe artifacts) based on SRS images[14]. **b** The intensity attenuation of mask is added to specific regions to simulate the stripe and artifacts of the stitched image. Scale bar: 1mm.

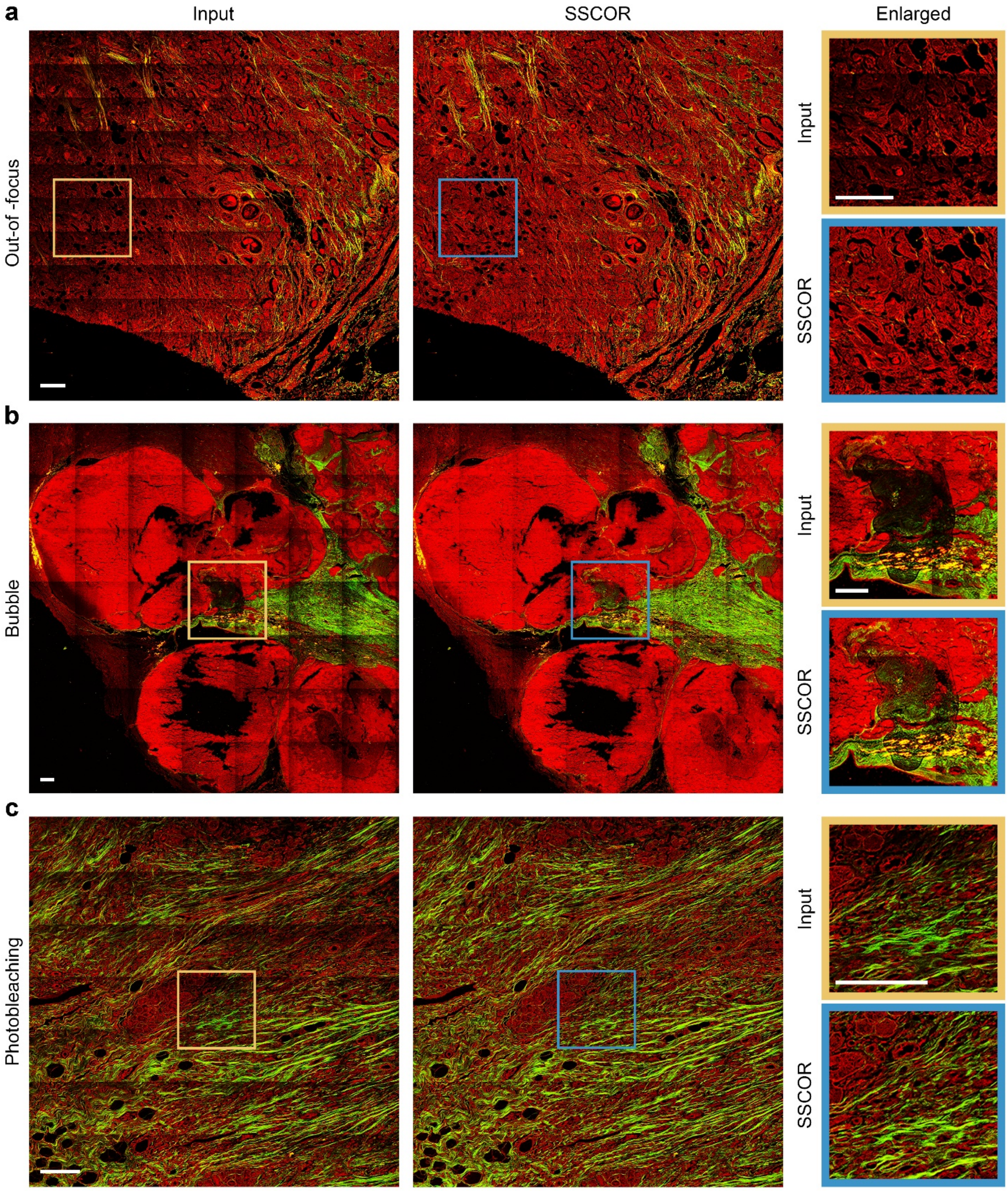


**Supplementary Figure 5. Restoration results of real artifacts in the MPM images. a** Compared with the input image, SSCOR can restore out-of-focus areas in the real MPM image of breast cancer while removing stripes. **b** SSCOR recovers the original tissue signal obscured by bubble-like artifacts (yellow box) in MPM images of cerebral cavernous malformations. **c** In imaging experiments, repeated scanning on the same position is prone to photobleaching artifacts (yellow box). SSCOR restores the photobleaching signal and correct stripe in the MPM image of breast cancer. Scale bars: 200 μm.

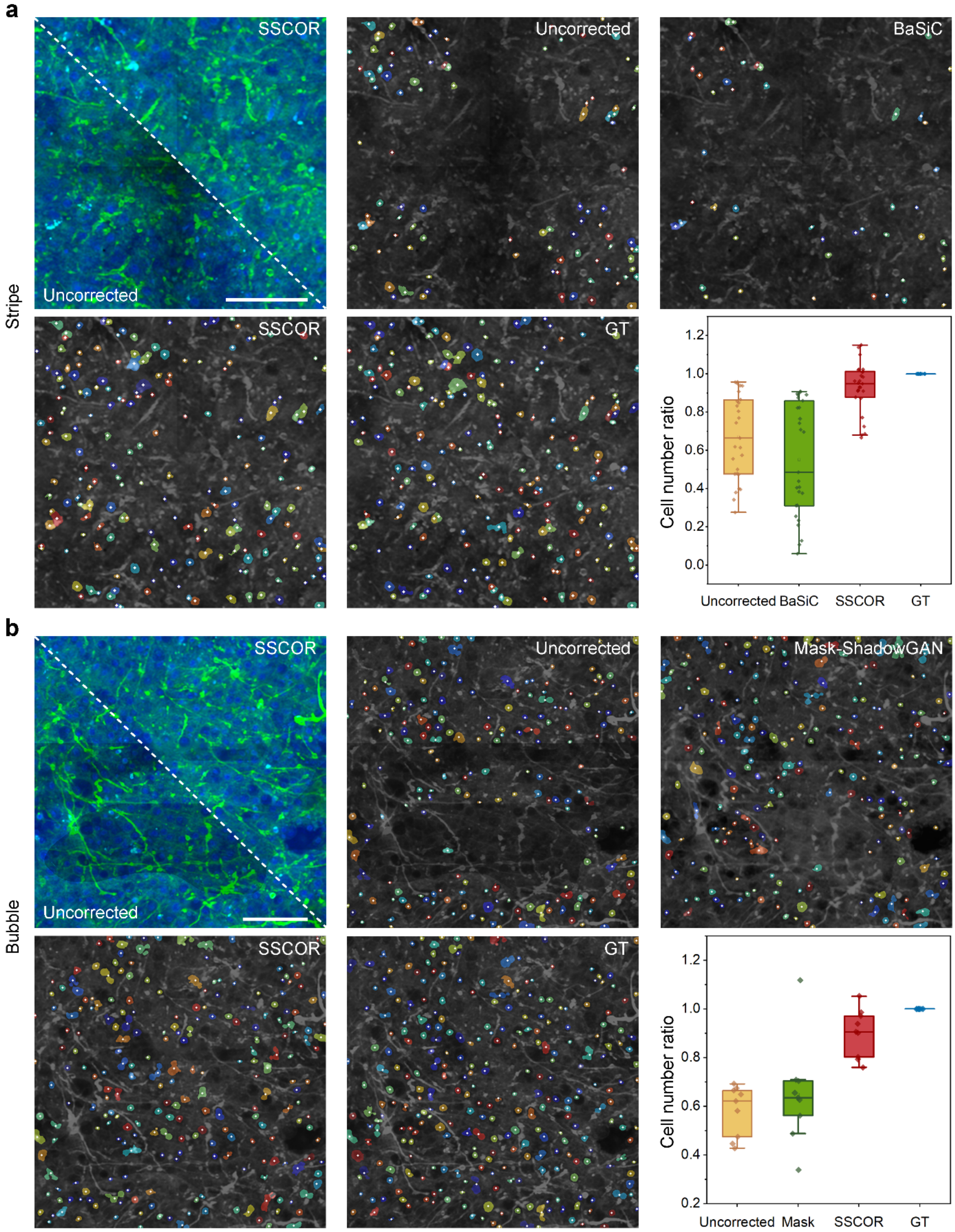


**Supplementary Figure 6. Cell counting results on bubble artifact and stripe images. a** and **b** Comparison of SSCOR-corrected images with uncorrected images, the automatic cell counting results of SSCOR are more consistent with ground-truth (GT) image. The original SRS image[14] before synthesis is considered as GT image. The cell number ratio quantitatively demonstrates the consistency of SSCOR-corrected and GT images ($n_{Stripe}$=27 and $n_{Bubble}$=8; $n$ represents the quantity of regions of interest (ROIs) in bubble-like artifact or stripe images). The white dots represent the location of cells, and the color-coded markers represent the size of cells. The cell number ratio is calculated by the ratio of cell numbers in the uncorrected/corrected image to the GT images. Scale bars: 100 μm.

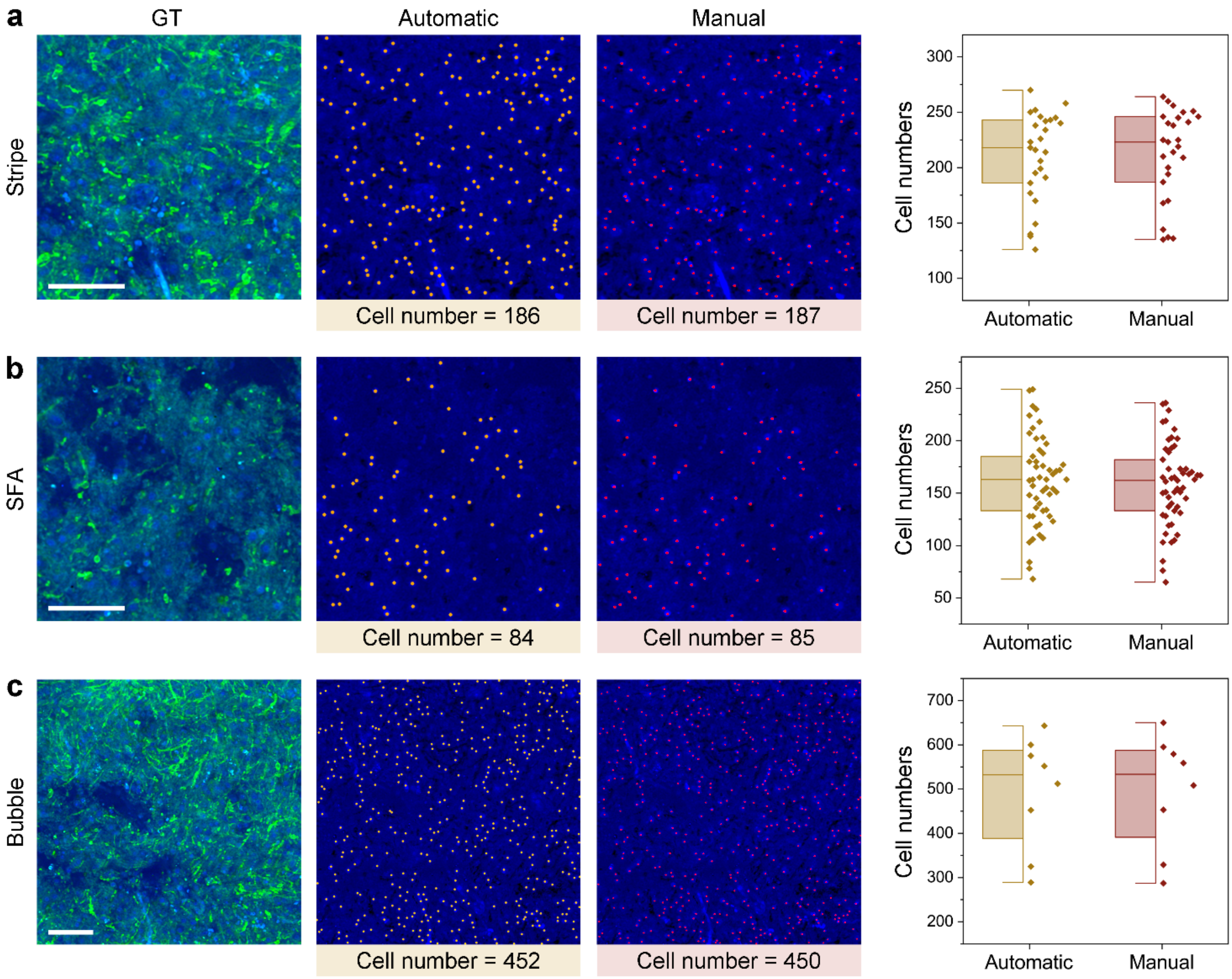


**Supplementary Figure 7. Accuracy of automatic counting compared with manual counting. a, b,** and **c** The comparison examples of automatic and manual cell counting on scanning and bubble artifacts and stripe images. The automatic cell counting[15] is performed on blue channel of SRS images[14]. The dots indicate the location of the cells. The quantitative results of cell numbers validate the accuracy of automatic counting ($n_{Stripe}$=27, $n_{SFA}$=53, and $n_{Bubble}$=8; $n$ represents the quantity of ROIs in artifacts or stripe images). Scale bars: 100 μm.

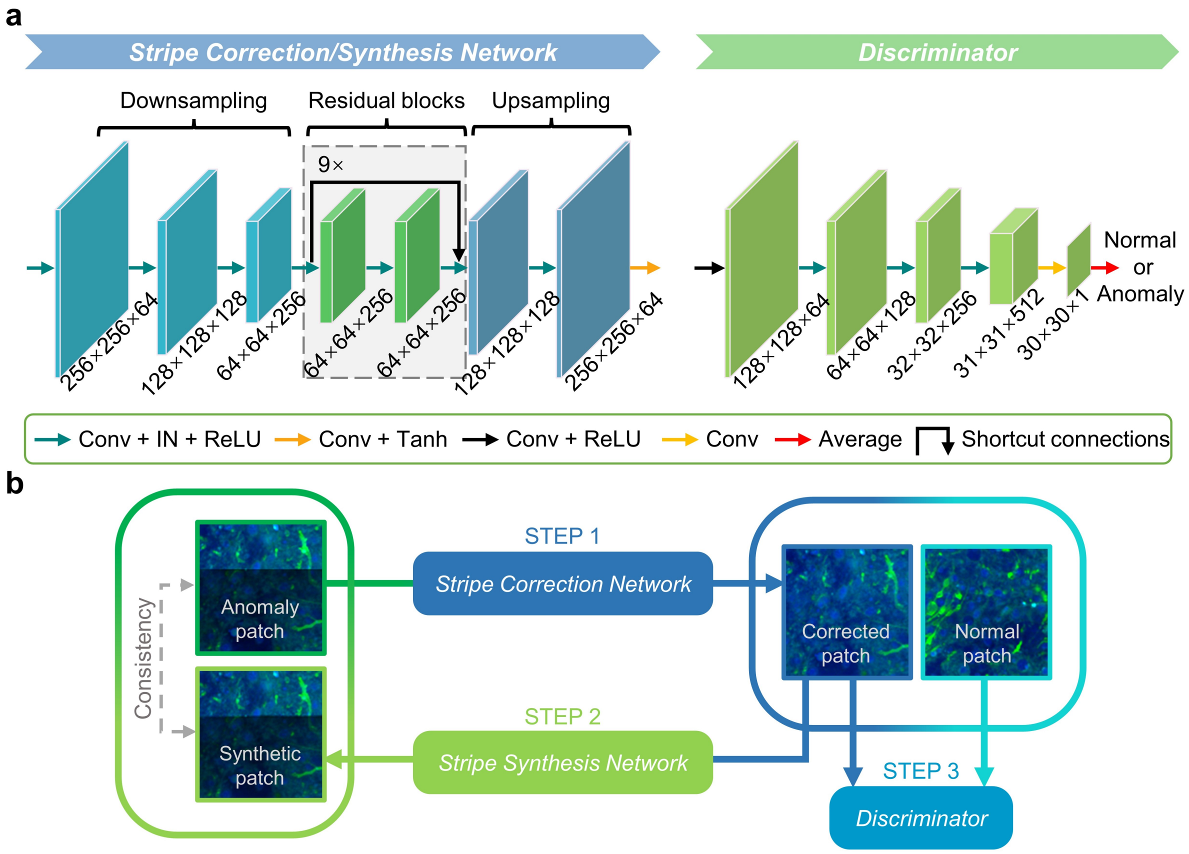


**Supplementary Figure 8. Visualization of network architecture. a** The stripe correction network consists of downsampling layers, intermediate layers, and upsampling layers. The discriminator network consists of five convolutional layers. **b** Diagram shows the image data flows during the adversarial self-training stage of SSCOR.

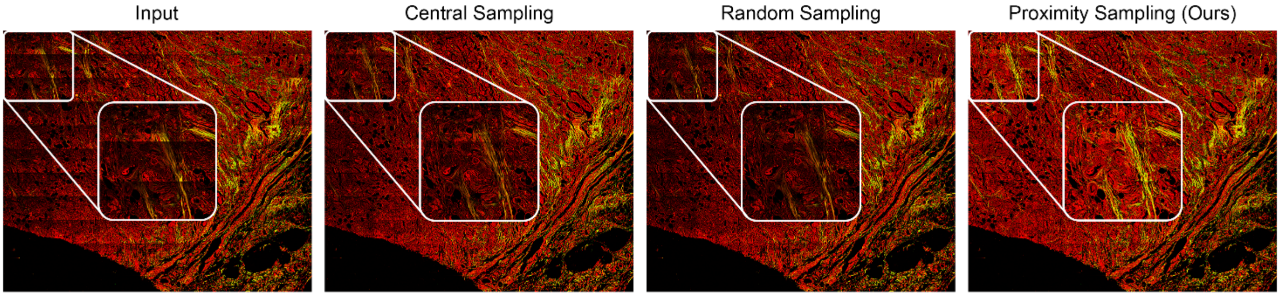


**Supplementary Figure 9.** Ablation study on the sampling method. Comparison of correction results among proximity sampling, central sampling and random sampling.

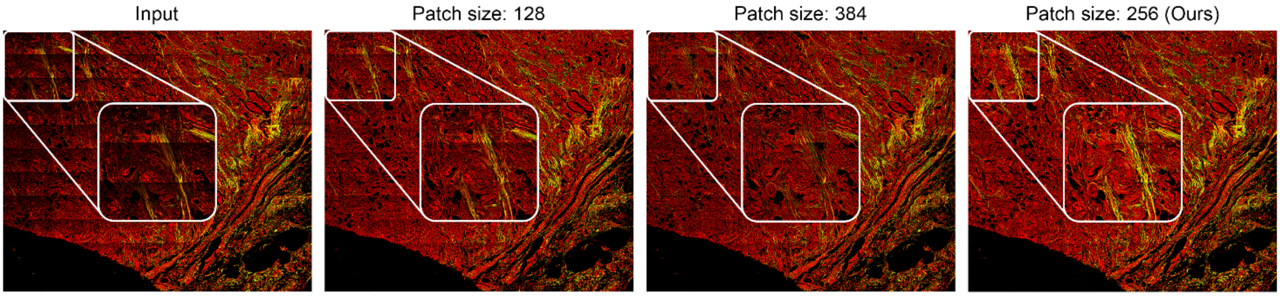


**Supplementary Figure 10.** Ablation study on the size of sampled patches. Setting different patch sizes for quantitative comparison.

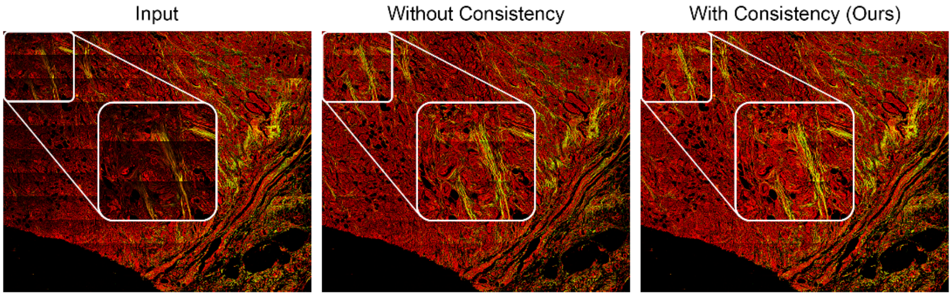


**Supplementary Figure 11.** Ablation study on the training scheme. The experiments are performed to access the importance of consistency constrain for stripe correction.

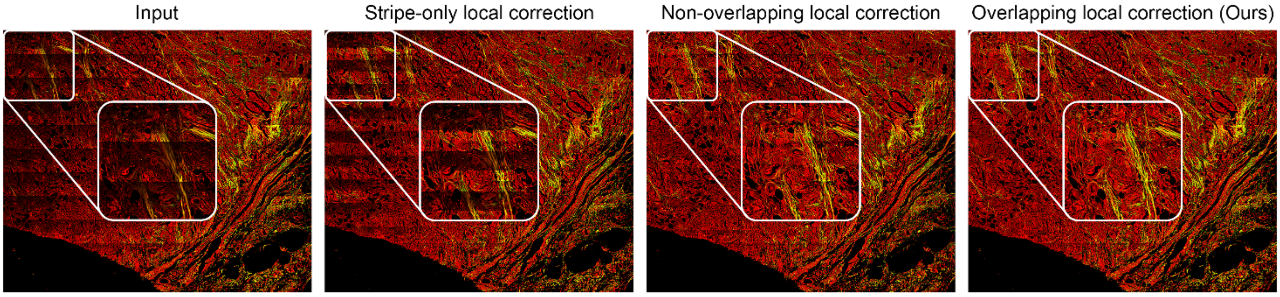


**Supplementary Figure 12.** Ablation study on the correction method. Correcting the stitched images with stripes in different methods to intuitively compare the correction quality.

### Supplementary Tables

#### Supplementary Table 1. Requirements of different methods for training images

| Method | Requirements for Image Training | | | | |
|---|---|---|---|---|---|
| | a 6348×5376 pixels stitched image with 13×11 tiles | | a 2881×2872 pixels stitched image with 3×3 tiles | | Image Format |
| | Sample Number | Sample Size (pixel) | Sample Number | Sample Size (pixel) | |
| ZEN[12] | 143 tiles | 512×512 | 9 tiles | 1024×1024 | .czi |
| BaSiC[11]<br>CIDRE[10] | | | | | .tif |
| Neighbor2Neighbor[7]<br>ZeroDCE[2] | 633 patches | 256×256 | 166 patches | 256×256 | .tif |
| Mask-ShadowGAN[5]<br>SSCOR | 633 unpaired patches | 256×256 | 166 unpaired patches | 256×256 | .tif |

#### Supplementary Table 2. Comparison of the microscopic datasets

| Stripes/Artifacts types | | Dataset | Specimen | Number | Resolution (pixel) | Microscope | Detector /Sensor | Objective | Immersion medium | Light Source | |
|---|---|---|---|---|---|---|---|---|---|---|---|
| | | | | | | | | | | Type | Detail |
| Stripes | Non-uniform | MPM | Breast cancer | 10 | 3430×3430, 8-bit | Zeiss LSM 880 | PMT | 20× 0.8 NA | Air | Coherent Laser | Chameleon Ultra Ti: Sapphire Femtosecond Excitation wavelength (810 nm) Average laser power (30 mW) |
| | | | Cerebral vascular malformation | 1 | 5550×5550, 8-bit | | | | | | |
| | Grid | | Liver cancer | 4 | 4403×4403, 8-bit | | | | | | |
| | Oblique | Fluorescence[13] | Mouse brain | 5 | 4171×3736, 8-bit | Hamamatsu L10387 | TDI | 20× 0.75 NA | Air | NanoZoomer Mercury lamp | LX2000 Ultrahigh-pressure 200 W |
| | | | | | 4346×3080, 8-bit | | | | | | |
| Artifacts | Out-of-focus | SRS[14] | Glioblastoma | 2 | 7350×5390, 8-bit | Olympus FV300 | CCD | 60× 1.2 NA | Water | APE GmbH Laser | picoEmerald Tunable Two-Color Source Picosecond Optimal Raman shifts (2973, 2921, and 2851 $cm^{-1}$) |
| | Bubble | | | | 2838×3892, 8-bit | | | | | | |
| | SFA | | | | | | | | | | |

MPM, multiphoton microscopy; PMT, photomultiplier; NA, numerical aperture; TDI, time delay integration; SRS, Stimulated Raman scattering; CCD, charge coupled device.
